# Caudal counter-represses Hunchback to regulate *even-skipped* stripe 2 expression in Drosophila embryos

**DOI:** 10.1101/226373

**Authors:** Ben J. Vincent, Max V. Staller, Francheska Lopez-Rivera, Meghan D.J. Bragdon, Zeba Wunderlich, Javier Estrada, Angela H. DePace

**Affiliations:** Department of Systems Biology, Harvard Medical School, Boston, MA 02115, USA; Present address: Department of Biological Sciences, University of Pittsburgh, Pittsburgh, PA 15260, USA; Present address: Department of Genetics and Edison Family Center for Genome Sciences & Systems Biology, Washington University School of Medicine in St. Louis, St. Louis, MO 63110, USA; Present address: Department of Biomedical Engineering and Biological Design Center, Boston University, Boston, MA 02215, USA; Present address: Department of Developmental and Cell Biology, University of California, Irvine, Irvine, CA 92697, USA

## Abstract

Hunchback is a bifunctional transcription factor that can activate and repress gene expression in Drosophila development. We investigated the regulatory DNA sequence features that control Hunchback function by perturbing enhancers for one of its target genes, *even-skipped*. While Hunchback directly represses the *eve* stripe 3+7 enhancer, we found that in the *eve* stripe 2+7 enhancer, Hunchback repression is prevented by Caudal binding—this relationship is called counter-repression. We found evidence that this relationship is conserved by comparing predicted binding sites for Hunchback and Caudal across orthologous *eve* stripe 2 enhancers. These results alter the textbook view of *eve* stripe 2 regulation wherein Hb is depicted as a direct activator. Instead, to generate stripe 2, Hunchback repression must be counteracted by Caudal binding. We discuss the implications of this interaction for *eve* stripe 2 regulation and evolution.

## Introduction

Bifunctional transcription factors (TFs) that can activate or repress their target genes are critical in animal development (1, 2) and are associated with human disease (3, 4). The function of these TFs can depend on the context of the enhancer sequences they bind (5–9). To infer accurate regulatory networks in development from regulatory DNA, we must define the sequence features that control the activity of bifunctional TFs (10).

Hunchback (Hb) is a bifunctional TF that patterns the *Drosophila melanogaster* embryo (11); the Hunchback homolog Ikaros is critical in human hematopoesis (12). *hb* is a gap gene with many targets throughout Drosophila development including other gap genes (13), pair-rule genes (14–17), homeotic genes (18–20) and neuronal genes (21). Its bifunctional role in regulating the pair-rule gene *even-skipped* has been particularly well-studied. Classic experiments measured expression driven by wild-type and mutant versions of *eve* enhancers and found that Hb directly activates the minimal *eve* stripe 2 enhancer *(eve2min)* (14, 22, 23). Other studies examined endogenous*eve* expression in embryos by misexpressing *hb* (*sna::hb* embryo) along the ventral surface of the embryo and found that Hb represses *eve* stripes 3 and 7 (24). Qualitative measurements of mutated versions of the *eve* stripe 3+7 enhancer (*eve3*+*7*)suggested that Hb repression of *eve3+7* is direct (15, 17, 25, 26).

Even after decades of study, the DNA sequence features that control Hb bifunctionality are not known. One hypothesis is that DNA-bound Hb dimers act as repressors while Hb monomers act as activators (dimerization hypothesis). The dimerization hypothesis is supported by computational work that predicts expression of *eve3*+*7* and the gap gene *Krüppel* (27, 28), as well as *in vitro* experiments identifying zinc finger domains in Hb and Ikaros that allow for dimerization (29). In another hypothesis, binding of a different TF converts Hb from a repressor to an activator (co-activation hypothesis). *In vivo* measurements of synthetic binding site arrays for Bicoid (Bcd) and Hb support the co-activation hypothesis (30), and Hb co-activation by Bcd has been incorporated into computational models of *eve* enhancer function (31–33). Recent models have also incorporated Hb co-activation by the TF Caudal (Cad) (34). Cad activates gap and pair-rule gene expression in the posterior of both long and short-band insects (35–38), and Cad homologs are critical in vertebrate development and human disease (39, 40). Though both hypotheses have been included in computational models of enhancer function (28, 34), neither has been experimentally tested.

Here we experimentally test the co-activation hypothesis by perturbing two enhancers that are active in the same cells but respond differently to Hb. *eve* stripe 7 is generated by two shadow enhancers; Hb represses *eve3*+*7* (24, 26) and activates *eve2*+*7*, an extended version of *eve2min* that also generates stripe 7 (41). Because both enhancers are active in the same nuclei, Hb function must be partially controlled by enhancer sequence. In this system, we can test the co-activation hypothesis by measuring the effects of perturbing both enhancers quantitatively and at cellular resolution (Fig. 1) (42–44).

**Fig 1.**
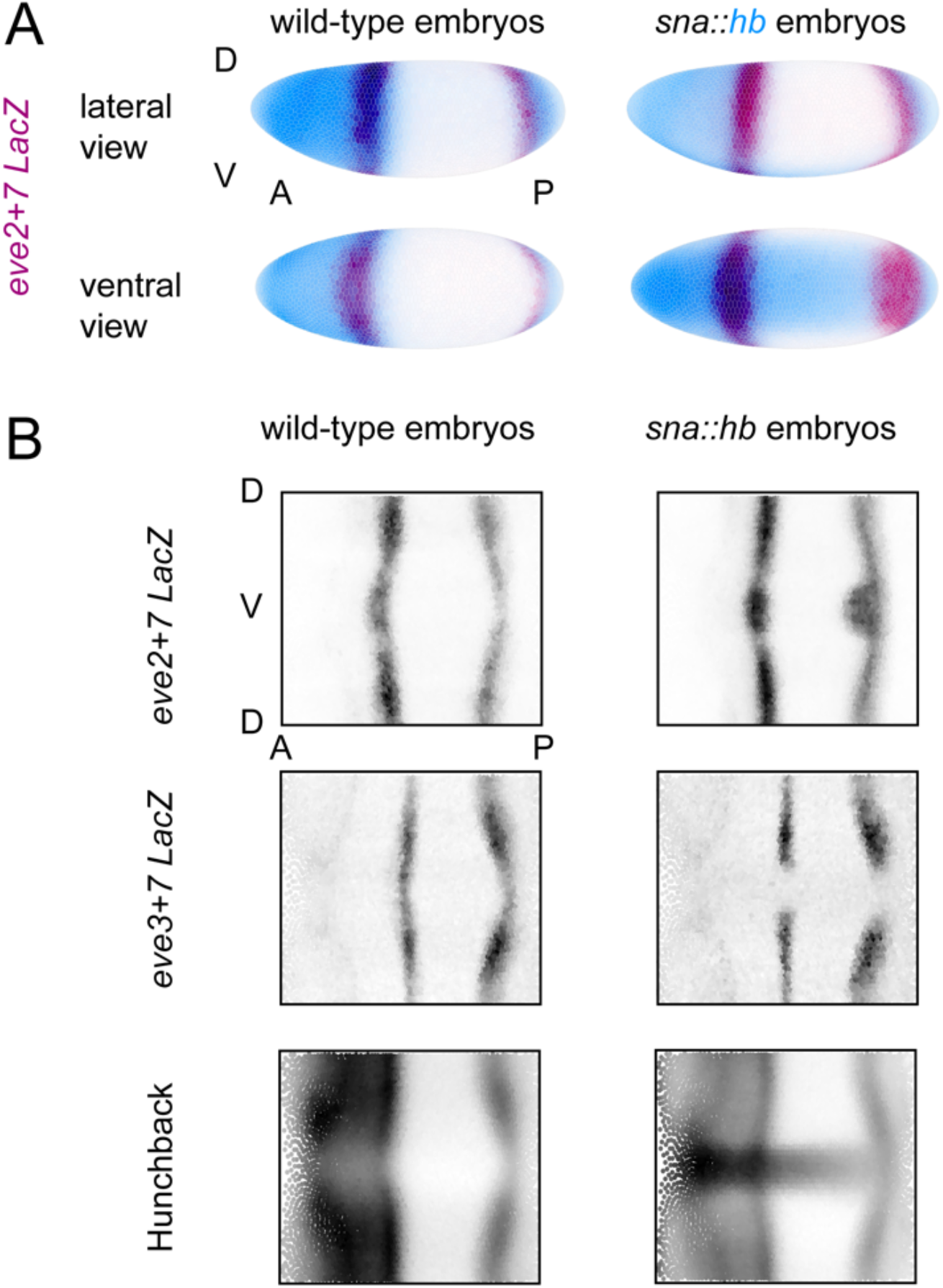
Quantitative gene expression data in wild-type and *sna::hb* embryos. (A) We created transgenic lines containing *lacZ* reporter constructs for *eve* enhancers and measured gene expression in wild-type embryos and embryos misexpressing ventral *hb* (*sna::hb*embryos). Here, we show visual renderings (66) of gene expression atlas data that average measurements from multiple embryos in time-point 4 (43). Dorsal (D) and ventral (V) surfaces are indicated for the lateral view, as are anterior (A) and posterior (P) positions. Left: *eve2*+*7 lacZ* expression (maroon) in wild-type embryos; Right: *eve2*+*7 lacZ* in *sna::hb* embryos. Hb protein is shown in blue. Individual nuclei are outlined, darker coloring indicates higher relative expression level. (B) To help visualize all relevant nuclei, we show 2-dimensional projections of expression data throughout the manuscript. Positions of individual nuclei along the dorsal-ventral axis are plotted as a function of position along the anterior-posterior axis; darker color indicates higher expression for each nucleus. Relative expression values are normalized to the maximum value and range from 0 to 1.

We find that Hb directly represses *eve3*+*7* and indirectly activates *eve2*+*7*. Indirect Hb activation occurs because Cad binding sites prevent Hb repression in *eve2*+*7*; this interaction appears to be evolutionarily conserved. This result alters the textbook description of *eve2min*, where Hb is depicted as a direct activator that synergizes with Bcd (45). Instead, Hb represses eve2, but this activity is masked by Cad, a direct regulator that counter-represses Hb. We discuss the possibility that additional activators may function as counter-repressors in development.

## Results

### Hb directly represses eve3+7

Mutating Hb binding sites in *eve3*+*7* previously showed that Hb defines the anterior border of stripe 3, but the effect of Hb on stripe 7 was unclear (26). The stripe 7 pattern driven by *eve3*+*7* retreats in response to ectopic Hb, but this effect could be indirect (41). To test whether Hb directly represses stripe 7 in the context of *eve3*+*7*, we removed predicted Hb binding sites from eve3+7 using the most current PWMs available, and measured expression using quantitative *in situ* hybridization in wild-type and *sna::hb* embryos (*eve3*+*7mutHb*, Fig. 2A). Some predicted Hb sites remain in *eve3*+*7mutHb* because they overlap with sites for other factors. We compared expression driven by this construct to expression driven by a wild-type version (Fig. 2) as well as the original mutant construct from Struffi et al. 2011 (Fig. S1) (26). All enhancers were cloned into the same reporter backbone and integrated into the same site in the Drosophila genome (see Materials and Methods). We display our quantitative data in multiple ways: a line trace of expression level versus anterior-posterior (AP) position along the lateral or ventral surface of the embryo; a 2-dimensional rendering of expression level in every cell in a single embryo or a gene expression atlas; or a plot of the differences in peak expression level between a lateral and a ventral line trace in individual *sna::hb* embryos to quantify the effect of *hb* misexpression.

**Fig. 2.**
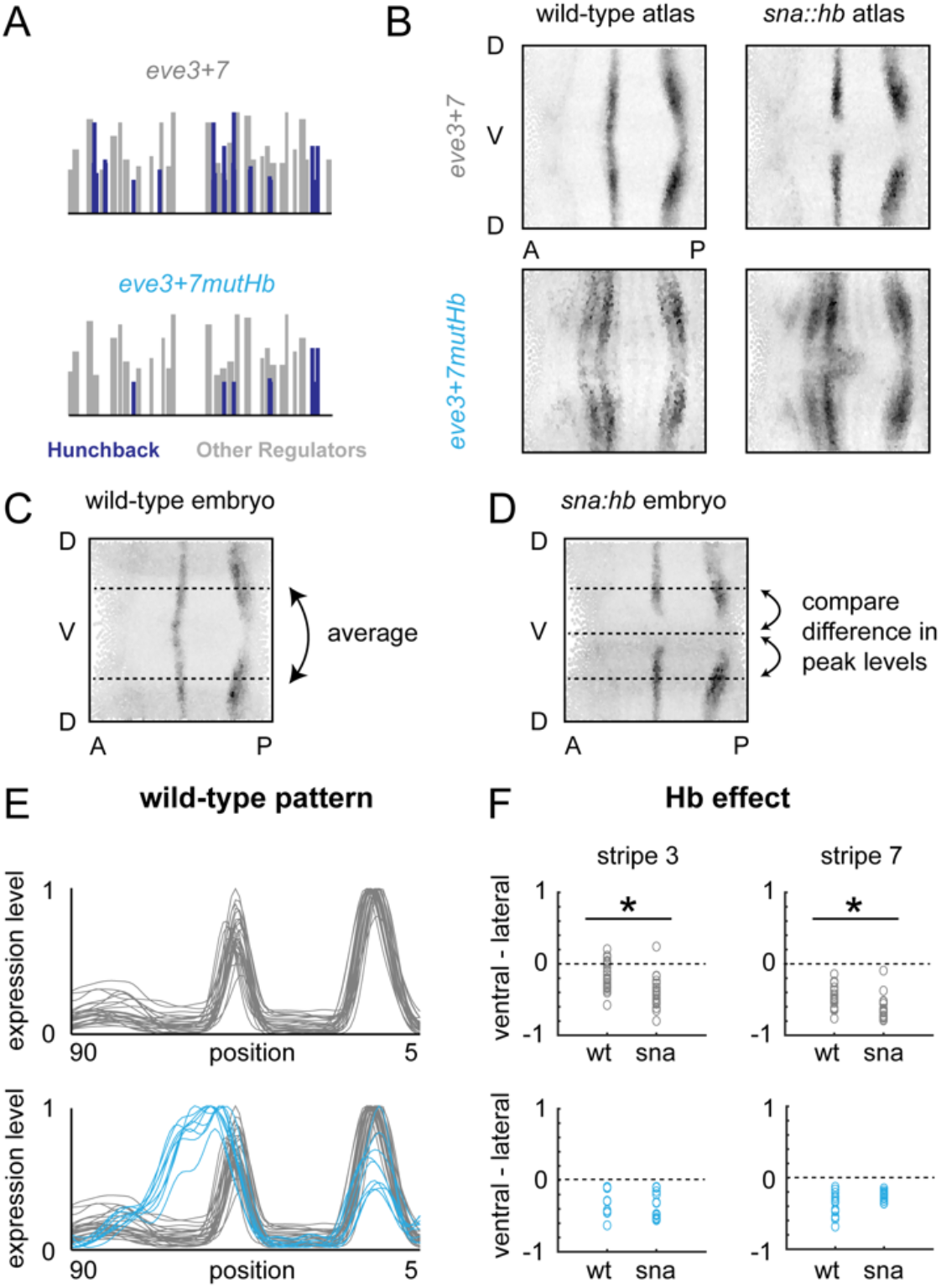
Hunchback directly represses *eve3*+*7*. (A) Predicted Hb binding sites in wild-type and mutant versions of *eve3*+*7* are depicted as vertical bars along the sequence where height is proportional to PATSER score (61). Hb sites are indicated in blue; other regulators are in grey. (B) 2D projections of atlas data for reporter constructs expressed in wild-type or *sna::hb* embryos is shown. Data is taken from timepoint 4. (C) 2D projection of a representative wild-type embryo expressing *eve3*+*7 lacZ*. We plot gene expression as a function of anterior-posterior position by averaging measurements from lateral strips in individual embryos and normalizing to the maximum value (see E). (D) 2D projection of a representative *sna::hb* embryo expressing *eve3*+*7 lacZ*. To quantify the effect of ventral Hb in individual embryos, we take average measurements from lateral strips, extract the local maxima corresponding to stripes 3 and 7, and subtract those values from the corresponding peaks from the ventral strip. We perform the same analysis in wild-type embryos to account for modulation along the dorsal ventral axis. A decrease in the ventral/lateral difference between wild-type and *sna::hb* embryos indicates Hb repression, while an increase indicates Hb activation (see F). (E) Lateral line traces from individual wild-type embryos containing *eve3*+*7* reporter constructs are shown (wild-type: grey, n= 26; mutant: blue, n = 9). Embryos are from all six timepoints in stage 5. (F) Differences between ventral and lateral stripe peaks are plotted for individual wild-type and *sna::hb* embryos in all 6 timepoints in stage 5. Top: wild-type *eve3*+*7* (wt: n = 26; *sna::hb:* n = 19); Bottom: *eve3*+*7mutHb* (wt: n = 9; *sna::hb:* n = 13). Asterisks indicate significant differences between wild-type and *sna::hb* embryos (p-value < 0.001, Mann-Whitney U test). Differences between wild-type and *sna::hb* embryos containing *eve3*+*7mutHb* were not significant (p-value > 0.4 for both stripes, Mann-Whitney U test).

Consistent with previous results (26), *eve3*+*7mutHb* drove an expression pattern where stripe 3 expanded anteriorly (Fig. 2B and E). In addition, the stripe 7 expression pattern driven by the *eve3+7mutHb* construct no longer retreats from ventral Hb in *sna::hb* embryos (Fig. 2B and F). These results confirm that Hb directly represses *eve3*+*7*.

### Hb indirectly activates stripe 7 in eve2+7

Previous experimental and computational work suggested that Hb directly activates *eve*stripe *2* (23, 31). We previously found that the stripe 7 pattern driven by *eve2*+*7* bulges in *sna::hb* embryos, and thus hypothesized that Hb directly activates *eve2*+*7* (41). Here, we tested whether Hb directly activates stripe 7 in *eve2*+*7* by mutating predicted Hb binding sites in *eve2*+*7* (*eve2*+*7mutHb*, Fig. 3A) and measuring gene expression in wild-type and mutant embryos.

**Fig. 3.**
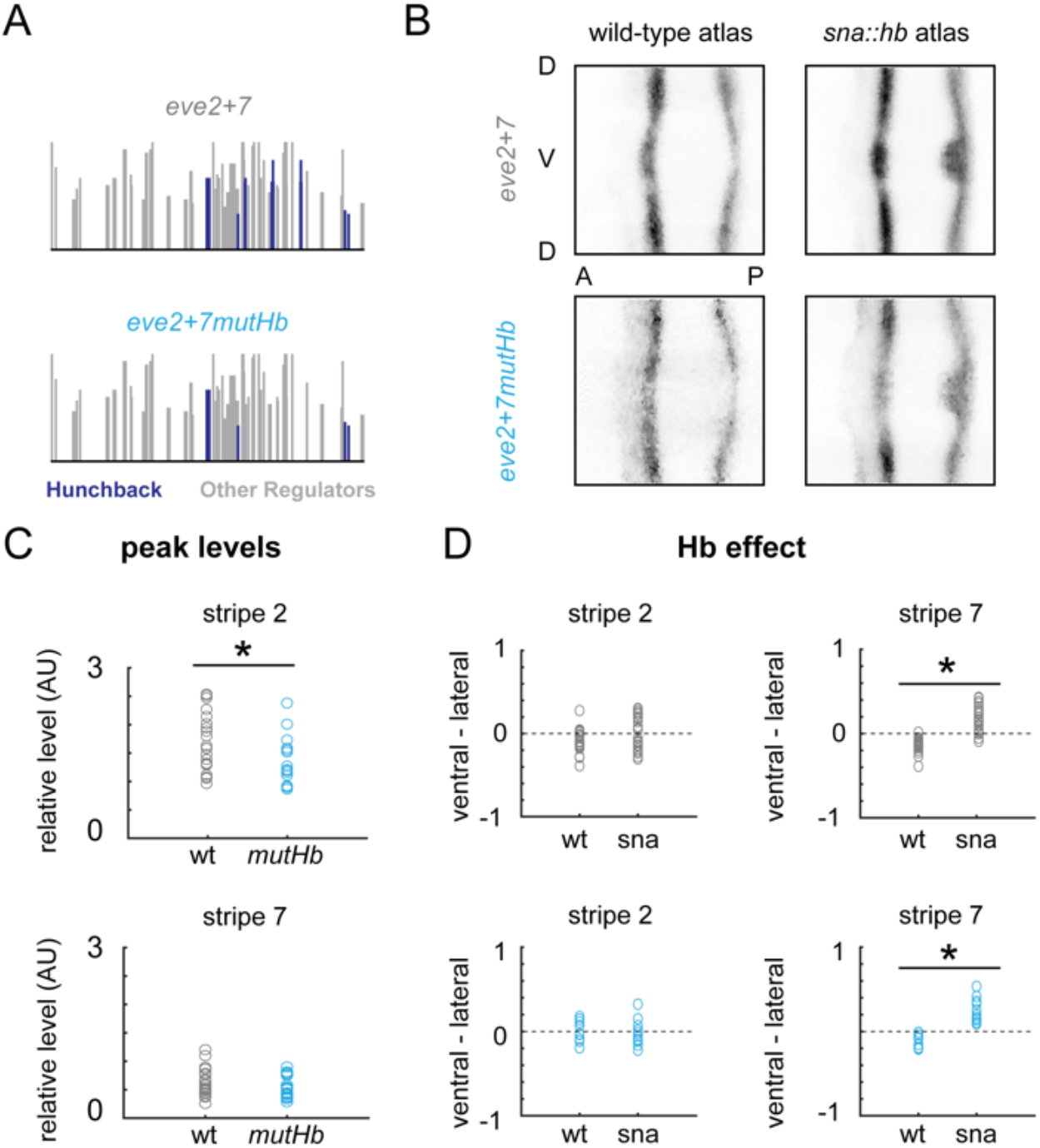
Hunchback indirectly activates *eve2+7*. (A) Predicted Hb binding sites in wild-type and mutant versions of *eve2+7*. Sites are displayed as in Figure 2. (B) 2D projections of atlas data for reporter constructs expressed in wild-type or *sna::hb* embryos. Data is taken from time point 4. (C) Peak stripe expression levels for individual embryos from timepoints 2-4 containing wild-type (grey, n = 23) and mutant (blue, n = 20) versions of *eve2*+*7*were measured by normalizing *lacZ* expression levels using a *huckebein* co-stain (44) and extracting local maxima from lateral line traces. Asterisks indicate significant differences (p-value < 0.05, Mann-Whitney U test). (D) Differences between ventral and lateral stripe peaks are plotted for individual wild-type and *sna::hb* embryos in all 6 timepoints in stage 5. Top: wild-type *eve2*+*7* (wt: n = 22; *sna::hb:* n = 26); Bottom: *eve2*+*7 mut Hb* (wt: n = 9; *sna::hb:* n = 11). Asterisks indicate significant differences between wild-type and *sna::hb* embryos (p-value < 0.001, Mann-Whitney U test).

We were surprised to find that mutating Hb binding sites had no effect on the spatial pattern driven by *eve2*+*7* in either genetic background (Fig. 3). We tested for quantitative effects of removing Hb binding sites by measuring expression levels using a co-stain method (44). We found that *eve2*+*7mutHb* drove slightly lower expression levels in stripe 2 (p-value = 0.022, Mann-Whitney U test), but stripe 7 levels were indistinguishable between *eve2*+*7* and *eve2*+*7mutHb* (p-value = 0.304, Mann-Whitney U test) (Fig 3C). Given the subtle effects that Hb binding site mutations had in *eve2*+*7*, we examined the behavior of other *eve* stripe 2 enhancer fragments in the literature. When we mutated all three footprinted binding sites in a fragment used in comparative studies (46), we observed a significant decrease in stripe 2 expression level (Fig S2). However, we observed no significant decrease in expression level as a result of mutating the footprinted Hb binding site in *eve2min* (Fig. S2), despite contradictory evidence from previous qualitative work (23, 47). These data indicate that Hb binding site mutations in *eve* stripe 2 enhancers have small, context dependent effects on gene expression. They further suggest that while Hb may be a weak activator of stripe 2, it does not influence stripe 7 expression at endogenous Hb levels in either *eve3*+*7* or *eve2+7*.

### Caudal binding prevents Hb repression of eve2+7

Our data indicate that Hb directly represses *eve3*+*7* but has little effect in *eve2*+*7*. In this system, the co-activation hypothesis makes a strong prediction: mutating Cad sites in *eve2*+*7*should convert Hb from an activator to a repressor of that sequence. We tested this hypothesis by mutating predicted Cad binding sites in *eve2*+*7* (*eve2*+*7mutCad*, Fig. 4A), and measuring gene expression in wild-type and sna::hb embryos. These mutations abolished stripe 2 expression completely and caused an anterior expansion of stripe 7 (Fig. 4B). We hypothesize that this anterior expansion of stripe 7 was caused by unintended mutations in binding sites for the repressor Giant (Gt) (see Discussion). In *sna::hb* embryos, removing Cad binding sites causes the stripe 7 pattern to retreat (Fig. 4B and D), confirming that without Cad, Hb behaves as a repressor in this sequence. To confirm that this effect was due to direct repression by Hb, we mutated both Hb and Cad binding sites in *eve2*+*7* (*eve2*+*7mutCadmutHb*). *eve2*+*7mutCadmutHb* drives a stripe 2 pattern that expanded to the anterior compared to WT *eve2*+*7* (Fig. 4B and C); we hypothesize this effect is also due to unintended mutations in Gt binding sites (see Discussion). Most importantly, these additional Hb mutations abolished repression by ventral Hb (Fig. 4). These results therefore support a variation of the co-activation hypothesis: Cad prevents Hb from directly repressing *eve2*+*7*.

**Fig. 4.**
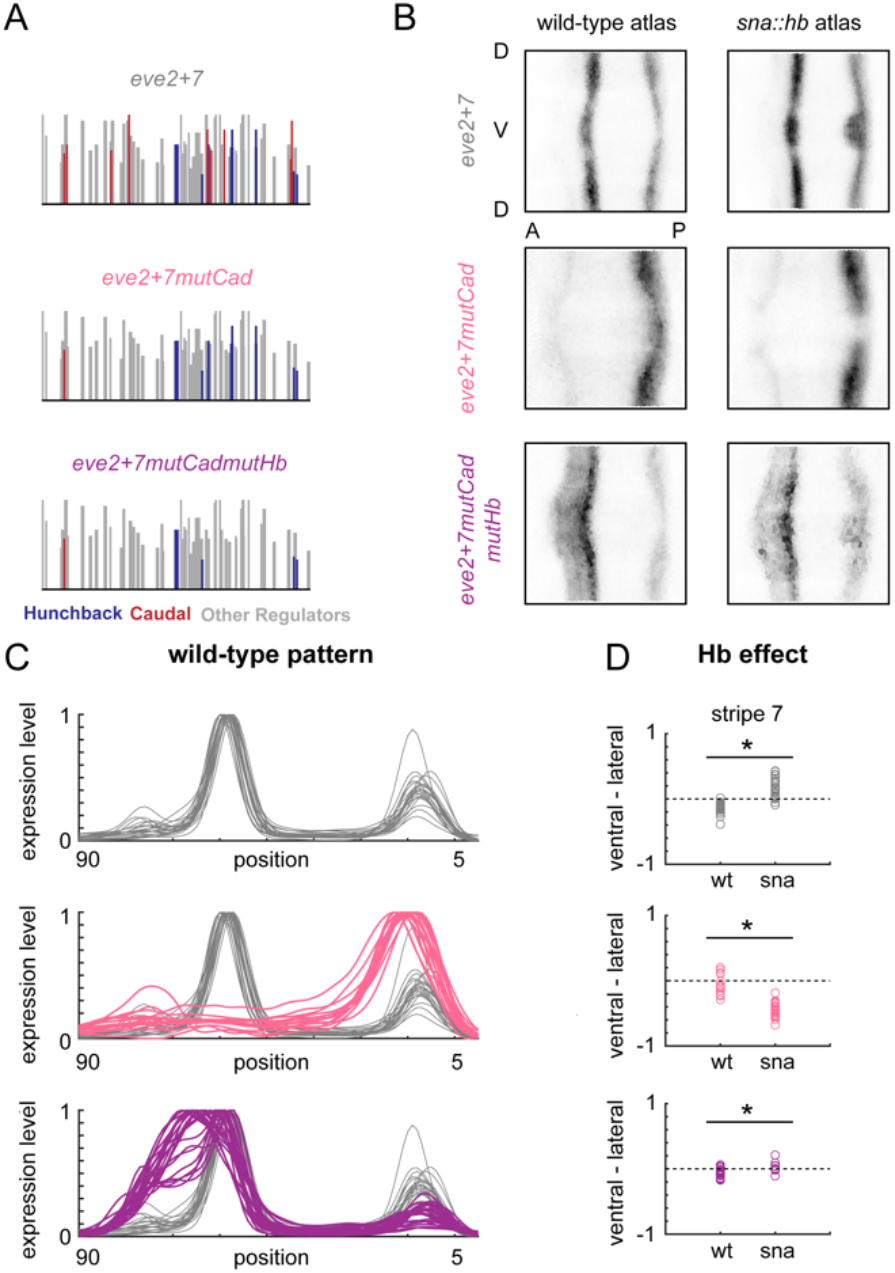
Caudal counter-represses Hunchback in *eve2*+*7*. (A) Predicted Cad (red) and Hb (blue) binding sites in wild-type and mutant versions of *eve2*+*7* are shown as in Figure 2. (B) 2D projections of atlas data for reporter constructs expressed in wild-type or *sna::hb* embryos. Data is taken from time point 4. (C) Lateral line traces from individual wild-type embryos containing *eve2*+*7* reporters are shown. Embryos are from all six timepoints in stage 5. (D) Differences between ventral and lateral stripe peaks are plotted for individual wild-type and *sna::hb* embryos in all 6 timepoints in stage 5. Top: wild-type *eve2*+*7* (wt: n = 22; sna: n = 26); Middle: *eve2*+*7mutCad*(wt: n = 15; sna: n = 24); Bottom: *eve2*+*7mutCadmutHb* (wt: n = 22; sna: n = 9). Asterisks indicate significant differences between wild-type and *sna::hb* embryos (p-value < 0.001, Mann-Whitney U test).

### Caudal binding sites co-evolve with Hunchback binding sites in eve2+7

Our results reveal that Cad sites in *eve2*+*7* ensure that Hb does not turn off *eve* stripe 2. We therefore hypothesized that the balance between Hb and Cad binding sites is under strong selective constraint, which would result in Hb and Cad binding sites co-evolving in *eve* stripe 2 orthologs. To test this hypothesis, we calculated binding site enrichment scores for Cad and Hb (48) in orthologous *eve* stripe *2* enhancers previously identified in Drosophila and Sepsid genomes (49). We found that Cad and Hb enrichment scores were significantly correlated in orthologous *eve* stripe 2 enhancers (Fig. 5A). In contrast, we did not observe any correlation between Hb and Cad binding sites in orthologous *eve3*+*7* enhancers when using a similar threshold for scoring predicted binding sites (Fig. 5A). This result is notable since mutating Cad binding sites in *eve3*+*7* decreases expression of both stripes (Fig. S3), suggesting that Cad also directly activates *eve3*+*7*. We also used the same method to also test for co-evolution of Hb and Bcd in *eve2* and *eve3*+*7* orthologs, but we observed no significant correlation in either case (Fig. S4). Although we cannot rule out the possibility that this interaction also occurs in *eve3*+*7*(see Discussion), these results suggest that Cad counter-repression of Hb is a conserved feature of *eve* stripe 2 enhancers.

**Fig. 5.**
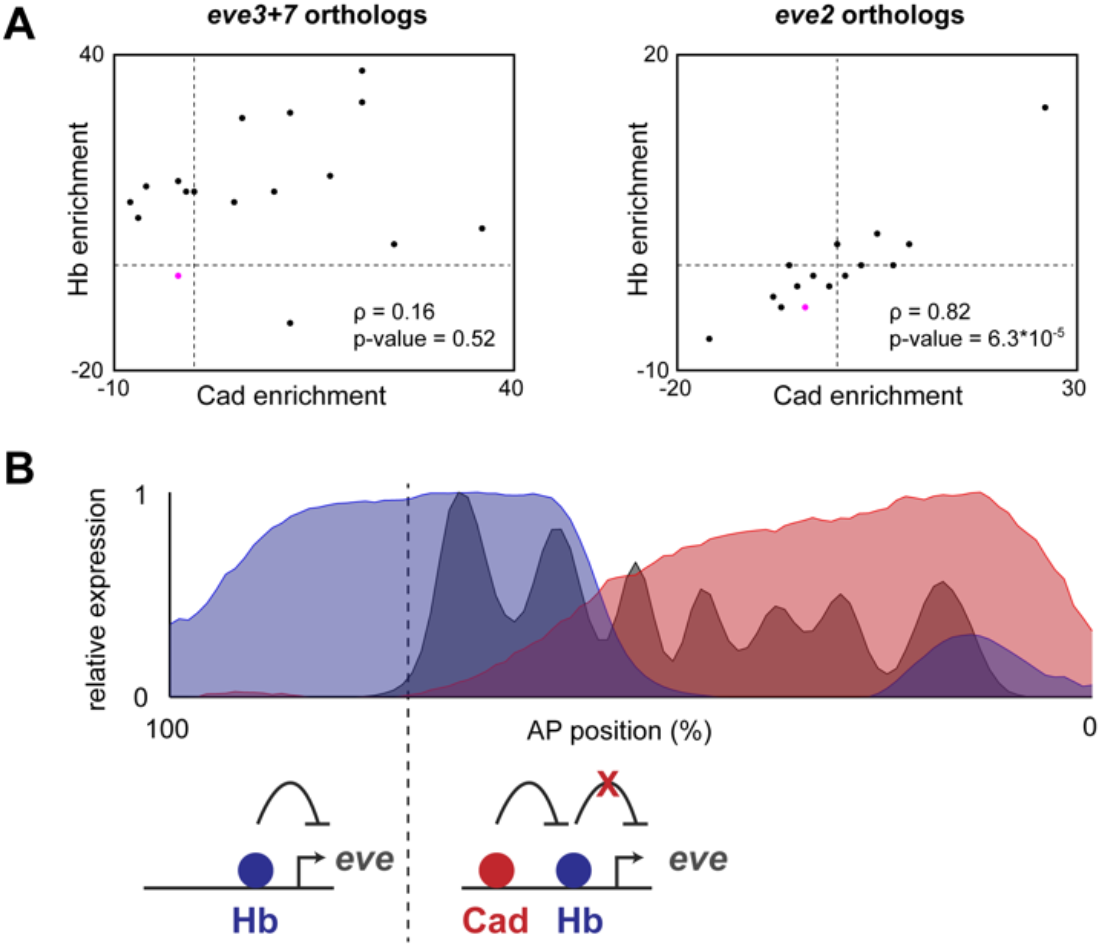
Caudal and Hunchback binding sites co-evolve in orthologous eve stripe 2 enhancers. (A) Enrichment scores for predicted Hb and Cad binding sites in orthologous *eve* stripe 2 and *eve3*+*7* sequences from different Drosophila and Sepsid species (49). Scores were calculated by comparing the number of predicted sites to an expected value calculated from the genomic background (48). Scores for *Drosophila me!anogaster* enhancers are indicated in magenta. Spearman correlation and p-value are displayed for each set of enhancers. (B) Summary of current findings. Top: levels of Hb protein (blue), Cad protein (red) and Eve protein (grey) are plotted as a function of anterior-posterior position. Data was taken from the FlyEx database (67). Bottom: cartoon indicating Hb function in different parts of the embryo. In the anterior, Cad levels are low, so Hb represses *eve2*+*7*. In the trunk, Cad binding to *eve2*+*7* prevents Hb repression.

## Discussion

The *eve* enhancers are often used to teach the fundamental principles of patterning (45, 50). Hunchback (Hb) is a key regulator of eve; it is thought to activate the *eve* stripe 2 enhancer in concert with Bicoid (Bcd), while it acts as a repressor at other enhancers, such as *eve3*+*7 (24, 26, 51)*. Here, we couple quantitative imaging and systematic perturbations of regulatory DNA and TF expression to uncover the DNA sequence features that control Hb activity. We show that Hb activity is controlled by a second TF, Caudal (Cad), and revise the classic description of *eve* stripe 2 regulation.

### Revising our picture of eve stripe 2 regulation

In the textbook picture of *eve* stripe 2 (45), Bcd and Hb directly activate the enhancer in the anterior of the embryo, while Gt and Kr directly repress it to carve out a single stripe of expression (14, 22, 23). However, attempts to reconstitute the minimal *eve* stripe 2 enhancer from its component binding sites have failed, which suggests that there is still more to learn about its regulatory logic (52). For example, these interactions alone cannot explain how *eve* stripe 2 is repressed in the anterior tip of the embryo. Two other mechanisms have been proposed to account for this: direct repression by Slp1 and downregulation of Bcd activity by Torso (53). Our results suggest an additional mechanism: Hb represses *eve2*+*7* in the anterior due to the absence of Cad (Fig 5B). In our data, an *eve* stripe 2+7 enhancer without Hb or Cad binding sites generates stripes 2 and 7 in the correct place with an anterior expansion of both stripes, which we hypothesize is due to mutations in Gt binding sites (see below). However, while an *eve2*+*7* enhancer lacking Hb and Cad sites could generate the correct pattern, its function could be easily compromised by the creation of Hb binding sites over evolutionary time. Regulation of *eve* stripe 2 by Hb and Cad therefores provide two different buffers against binding site mutations: Hb mediated repression in the anterior may allow for loss of other repressor binding sites (such as Slp1), and counter-repression between Cad and Hb guards against intrusion of Hb binding sites, which would turn off *eve* stripe 2 without Cad.

The correlation we observe in *eve* stripe 2 enhancers may reflect counter-repression in that enhancer specifically. However, this bioinformatic signature is not enough to rule out the possibility that Cad also counter-represses Hb in *eve3*+*7*. Sensitivity analyses indicate that the correlation of Hb and Cad binding sites in *eve* stripe *2* orthologs is highest when sites with lower PATSER scores are included, while in *eve3*+*7* orthologs we find that correlation values generally increase with PATSER threshold (Fig S5). These results may reflect differences in the constraints on the regulatory logic of *eve2*+*7* and *eve3*+*7*. Hb repression in *eve2*+*7* must be prevented or counteracted by Cad to allow for stripe 2 expression, whereas Hb repression in *eve3*+*7* is necessary for correct positioning of the stripes but may be modulated by Cad binding. Nonetheless, the co-evolutionary signature we observe in *eve* stripe *2* orthologs suggests that the functional interaction between Cad and Hb sites may be a conserved feature of *eve* stripe 2 regulation.

Bcd and Hb have been proposed to synergistically activate *eve* stripe 2 (14, 30), but our results do not support this hypothesis. Computational models that include Bcd/Hb synergy predict that Hb binding site mutations cause large decreases in expression (31). However, our data suggest only modest effects of Hb binding site mutations in accordance with qualitative data in the literature (23, 47). In addition, our results do not support the previous co-activation model for Bcd/Hb synergy in *eve2*+*7*; without Cad, Hb represses *eve2*+*7* even though Bcd binding sites are the same.

### Hb ‘bifunctionality’ is due to counter-repression

Our previous work indicated that Hb activates *eve2*+*7* and represses *eve3*+*7*. Because Hb was “known” to be an activator of *eve2min* and repressor of *eve3*+*7*, we hypothesized that these interactions were direct. Here, we find that Hb activation of *eve* stripe 7 is indirect. We now hypothesize that *sna::hb* causes retreat of Gt and subsequent anterior expansion of the stripe 7 pattern (41). Knirps forms the anterior boundary of stripe 7 through the *eve3*+*7* enhancer (26), while Gt has been proposed to form the anterior boundary of stripe 7 through *eve2*+*7* (31). Indeed, we believe that the anterior stripe 7 expansion we observe after mutating Cad sites in *eve2*+*7* is caused by unintended mutations in Gt binding sites. Predicted Gt sites overlap with predicted Cad sites in this sequence, and our Cad mutations disrupt a predicted Gt site that overlaps a footprinted Gt site in *eve2min* (Fig S6) (54). While these effects make it difficult to determine whether Cad directly activates *eve2+7*, they do not affect our conclusions concerning the influence of Cad on Hb function. Which TFs activate stripe 7 expression is less clear, though Cad and Zelda are obvious candidates shared by *eve2*+*7* and *eve3*+*7*. Overall, the conclusion of our previous paper—that *eve* stripe 7 shadow enhancers use different regulatory logic—still holds. Hb represses *eve3*+*7* but has little effect on *eve2*+*7* because of counter-repression. Furthermore, we hypothesize that the two enhancers use different repressors to set the anterior border, though this remains to be tested directly. We also previously showed that Hb activates and represses separate *Kr* shadow enhancers using the same *sna::hb* misexpression assay (48); it is thus possible that Cad also counter-represses Hb in the proximal *Kr* enhancer.

Cad may counter-repress Hb through any number of molecular mechanisms. For example, Cad may interfere with Hb via direct protein-protein interactions. The simplest idea is that when Cad is present, Hb does not bind. This isn’t supported by genome-wide chromatin immunoprecipitation (ChIP) data for both Hb and Cad; both are bound to *eve2*+*7* (55, 56). We note two caveats of this data. First, it is not spatially resolved, so it remains possible that Hb binding is affected in a subset of cells. Second, Cad may disrupt Hb binding quantitatively, enough to affect function but not to detect changes by ChIP. Another possibility is that direct protein-protein interactions between Cad and Hb interfere with Hb protein domains that execute its repressive function (21). Finally, interactions could occur indirectly through co-regulators, or mechanisms analogous to the effects of short-range repressors on nearby activators (57). Importantly, Cad is not the only counter-repressor in Drosophila development. Stat92E functions as a counter-repressor in the formation of the posterior spiracle, but its target repressor remains unknown (58). Different counter repressors may exert their effects through different molecular mechanisms.

We note that endogenous *eve* stripe 2 is expressed in *caudal* mutant embryos (38), which conflicts with our result that mutations in Cad binding sites in *eve2*+*7* abolish stripe 2 expression. This discrepancy may be explained in a number of ways. First, *hb* expression may be affected in a Cad mutant background. While *hb* expression is detectable in this background by qualitative *in situ* hybridization (38), *hb* expression levels may be lower, which may affect its capacity to repress stripe 2. Second, the piece of DNA that we tested in reporters may not contain all relevant DNA that contributes to *eve* stripe 2 expression. Indeed, in the endogenous *eve* locus, the DNA between *eve2*+*7* and *eve3*+*7* contains binding sites for many *eve*regulators, but is notably devoid of predicted Hb binding sites (41). It may therefore not be affected by Hb repression in the absence of Cad. Finally, the placement of the *eve2*+*7* enhancer adjacent to the promoter in our reporter constructs may allow Hb to exert a stronger repressive effect than in the endogenous locus. As discrepancies between *eve3*+*7* and the endogenous stripes revealed the influence *eve2*+*7* as a stripe 7 shadow enhancer (41), further dissection of stripe behavior in *cad* mutant embryos may yield additional insights into *eve* regulatory logic over the entire locus.

A long-term goal of studying gene regulation is predicting the output of a specific enhancer (10). Context-dependent function of TFs [e.g. (59)] is a huge hurdle to meeting this goal. Here we further elucidate counter-repression, one type of context dependence, and offer some strategies for uncovering it in regulatory DNA. First, single mutations of annotated activators or bifunctional TFs may be misleading. Analyzing the effects of single and double mutations together are necessary to classify activators, repressors, counter-repressors and the targets of counter-repression. For example, let’s say that you mutate binding sites for TF #1 and expression decreases. TF #1 may either be an activator or a counter-repressor. You now combine mutations in TF #1 with mutations in TF #2, and expression is restored. TF #2 may be a repressor or the target of counter-repression. Finally, you examine the the effect of mutating TF #2 alone. If TF #2 is a repressor, expression will increase, but if TF #2 is the target of counter-repression, there will be no effect. All three experiments are necessary to correctly classify both TF #1 and TF #2. High-throughput mutational studies both in reporters (9) and *in vivo* (60) may be able to systematically gather this type of data on combinatorial effects.

### Conclusion

By combining targeted genetic perturbations with quantitative expression measurements, we uncovered counter-repression as a key feature of *eve* stripe 2 regulation. This type of TF interaction, which converts Hb from a repressor to a “nonfunctional” regulator, may be useful for preventing activation in the anterior of the embryo, or for buffering against the effect of binding site introgression. In both cases, counter-repression increases the flexibility of *eve* stripe 2 regulatory logic, which may impact the plasticity of the enhancer over evolutionary time.

## Materials and Methods

### Binding site predictions and construct design

We used PATSER (61) to predict binding sites in *eve3*+*7* and *eve2*+*7* for blastoderm TFs. We used position weight matrices (PWMs) derived from bacterial 1-hybrid experiments for the following factors: Bicoid, Caudal, Dichaete, Stat92E, Hunchback, Krüppel, and Nubbin (62). We used other published PWMs for Giant, Knirps, and Tailless (63). Finally, we used a Zelda PWM from a personal communication with Christine Rushlow. Count matrices were converted into frequency matrices for use in PATSER using a pseudocount of 1. In designing binding site mutations, we predicted the effects on predicted sites above a p-value cutoff of 0.003.

### Fly work

All reporter constructs were cloned into the NotI/BglII insertion site in the pBϕY reporter plasmid (49) and integrated into the attP2 landing site using injection services provided by BestGene Inc (64). Successful transformants were homozygosed using the mini-white marker. We generated *sna::hb* embryos as described previously (24) using two copies of the *sna::hb* transgene, a generous gift from Steve Small.

### In situ hybridization

Embryos were collected, fixed in heptane and paraformaldehyde, stained using quantitative *in situ* hybridization, and imaged using 2-photon confocal microscopy as described previously (42, 48). For each embryo, we measure *fushi-tarazu* mRNA using an anti-digoxigenin horseradish peroxidase (HRP) antibody (Roche) and a coumarin-tyramide color reaction (PerkinElmer), as well as *lacZ* mRNA using an anti-2,4-dinitrophenyl HRP antibody (PerkinElmer) and a Cy3-tyramide color reaction (PerkinElmer). The *sna::hb* crossing scheme generates wild-type and mutant embryos in approximately equal proportions; we identified embryos misexpressing *hb* using the *ftz* expression pattern. For experiments involving a *hkb* costain, we stained embryos in the same batch, discarded outliers, and ensured that *hkb* levels and *lacZ* levels were significantly correlated as described previously (44, 48)

### Binding site overrepresentation analysis

Binding site enrichment was performed as described previously (48). To perform this analysis, we used PWMs from FlyFactorSurvey (65) with sequencing data collected on the SOLEXA platform. For our sensitivity analysis, we systematically varied the PATSER score cutoff used to identify relevant sites and calculated the effect on the Spearman correlation coefficient.

## Acknowledgements

The authors gratefully acknowledge Steve Small for the *sna::hb* flies, as well as Clarissa Scholes, Kelly Biette, and all members of the DePace Lab for helpful comments on the manuscript. This work was generously supported by the Albert J. Ryan Foundation (to B.J.V), the Harvard Herchel Smith Graduate Student Fellowship (to M.V.S.), the Jane Coffin Childs Memorial Fund for Medical Research (to Z.W.), the National Institutes of Health (U01GM103804, R21HD072481, and R01GM122928 to A.H.D., as well as K99/R00 HD73191 to Z.W.), the National Science Foundation (CAREER-iOS 1452557), the Giovanni Armenise Foundation (to A.H.D.), the Novartis Fellows Fund (to A.H.D.), and the McKenzie Family Charitable Trust (to A.H.D.).

---

**Fig. S1:**
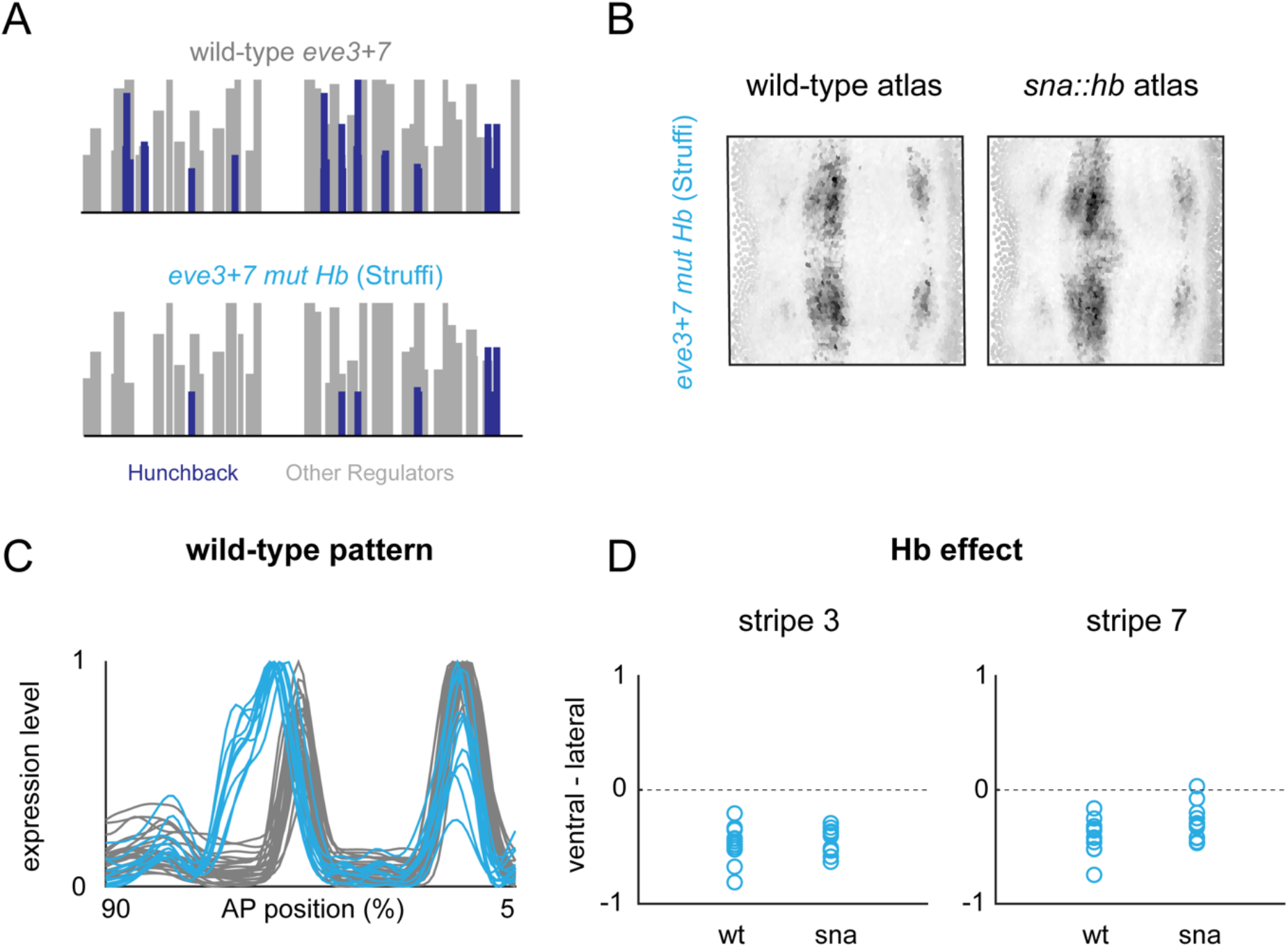
Quantitative effects of *eve3*+*7* mutations designed by Struffi et al. (A) Predicted Hb binding sites (cobalt blue) in *eve3*+*7* and *eve3+mutHb* plotted as in Figure 3.2. Sequence from Struffi et al. 2011. (B) 2D projections of atlas data for reporter constructs expressed in WT or *sna::hb* embryos. Data is taken from time point 4. Low-level anterior and posterior expression is due to an unused *hkb* co-stain. (C) Lateral line traces from individual WT embryos containing *eve3*+*7* reporter constructs (WT: grey, n= 26; *mutHb:* blue, n = 11). Each trace is normalized to its maximum value. Embryos are from all six time points in stage 5. (D) Differences in the maximum values of ventral and lateral line traces are plotted for individual WT and *sna::hb*embryos containing *eve3*+*7 mutHb i*n all 6 time points in stage 5. wt: n = 11; *sna::hb:* n = 12. Differences between WT and *sna::hb* embryos were not significant (Mann-Whitney U test, p-value > 0.1 for both stripes).

**Fig. S2:**
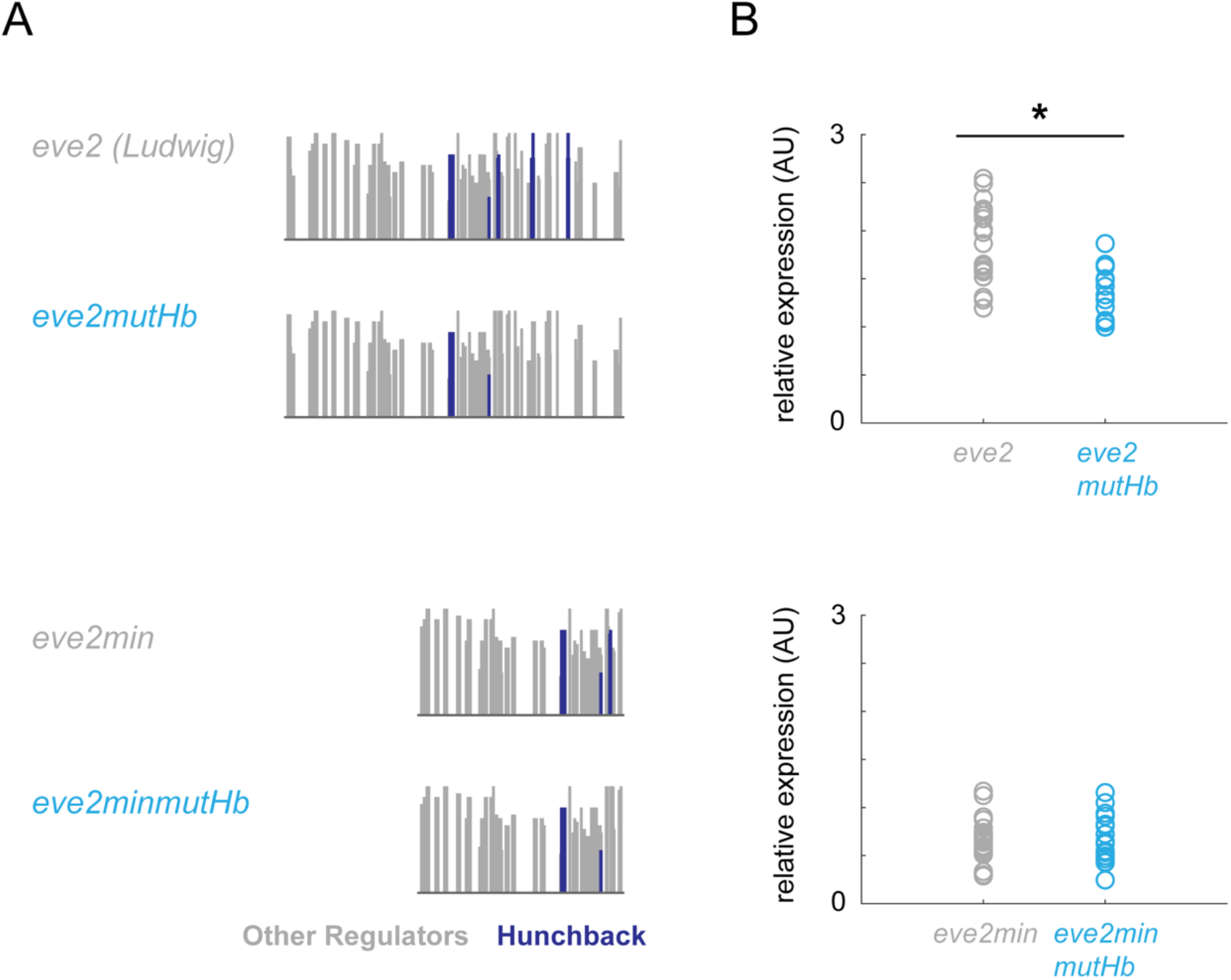
Hunchback mutations have different effects depending on enhancer context. (A) Predicted binding sites for Hb (cobalt blue) and other *eve2*+*7* regulators (grey) in different stripe 2 enhancer constructs (46,47). (B) Peak stripe 2 expression levels for individual embryos from time points 2-4 were measured using a *hkb* co-stain method (44). Asterisks indicate p-values < 0.05 (Mann-Whitney U test). Note that because each experiment was performed in separate hbridizations, comparisons can only be made between wild-type and mutated versions of the same enhancer, not between different stripe 2 enhancers.

**Fig. S3:**
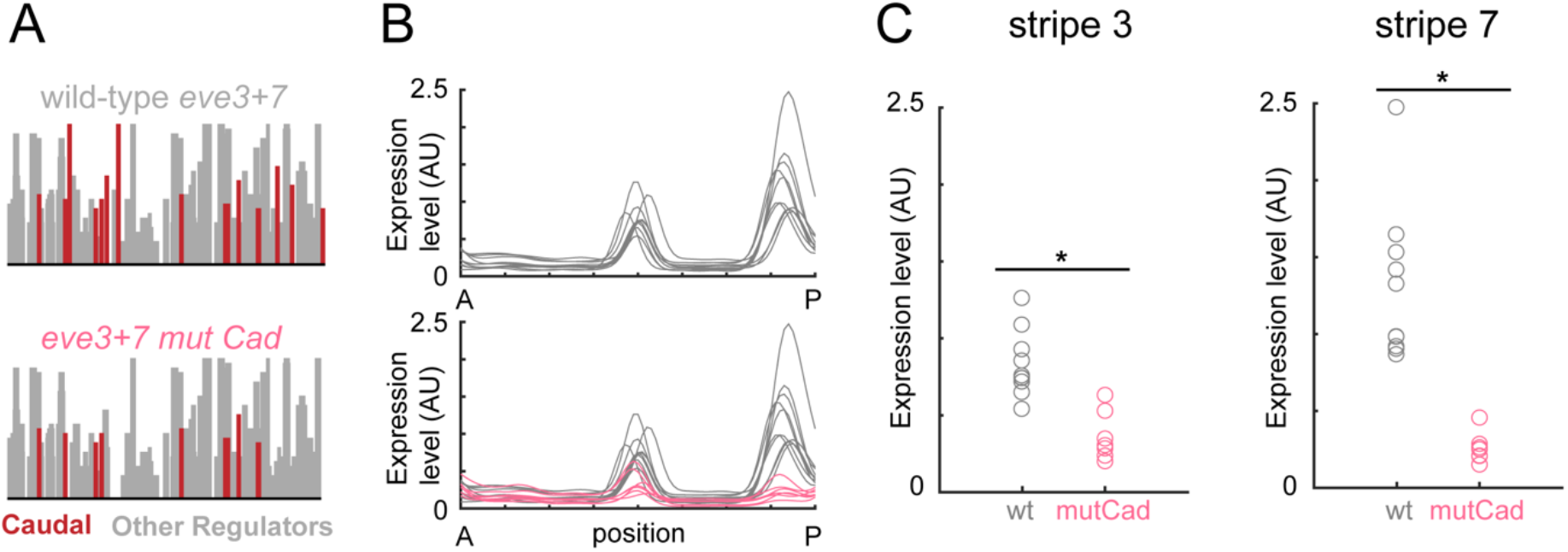
Caudal directly activates *eve3*+*7*. (A) Predicted Cad binding sites in *eve3*+*7* and *eve3*+*7mutCad*. Sites were predicted and displayed as described in previous figures. (B) Lateral line traces from individual wild-type embryos containing reporter constructs for *eve3*+*7* (grey, n = 11) and *eve3*+*7mutCad* (carmine, n = 7). Traces were normalized using a co-stain method (44); embryos are from time points 3 and 4. (C) Individual stripe peaks were found by taking local maxima from line traces in B. Asterisks indicate significant differences in stripe level (p-values < 0.001, Mann-Whitney U test).

**Fig. S4:**
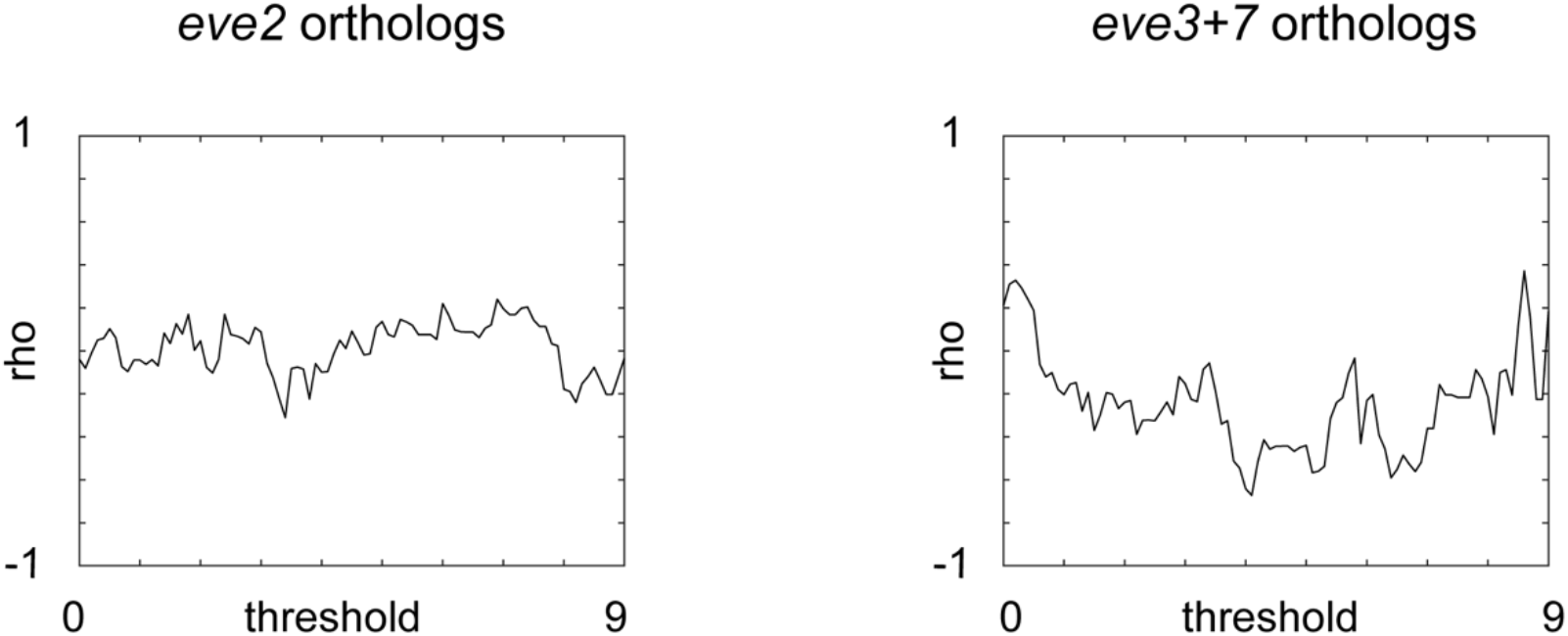
Sensitivity analyses for Bicoid and Hunchback enrichment correlations. (A) Spearman correlation for Bcd and Hb binding site enrichment is plotted as a function of binding site threshold for *eve2* and *eve3*+*7* orthologs. Enrichment for Bcd and Hb sites are not significantly correlated at any binding site threshold in either *eve2* or *eve3*+*7* orthologs.

**Fig. S5:**
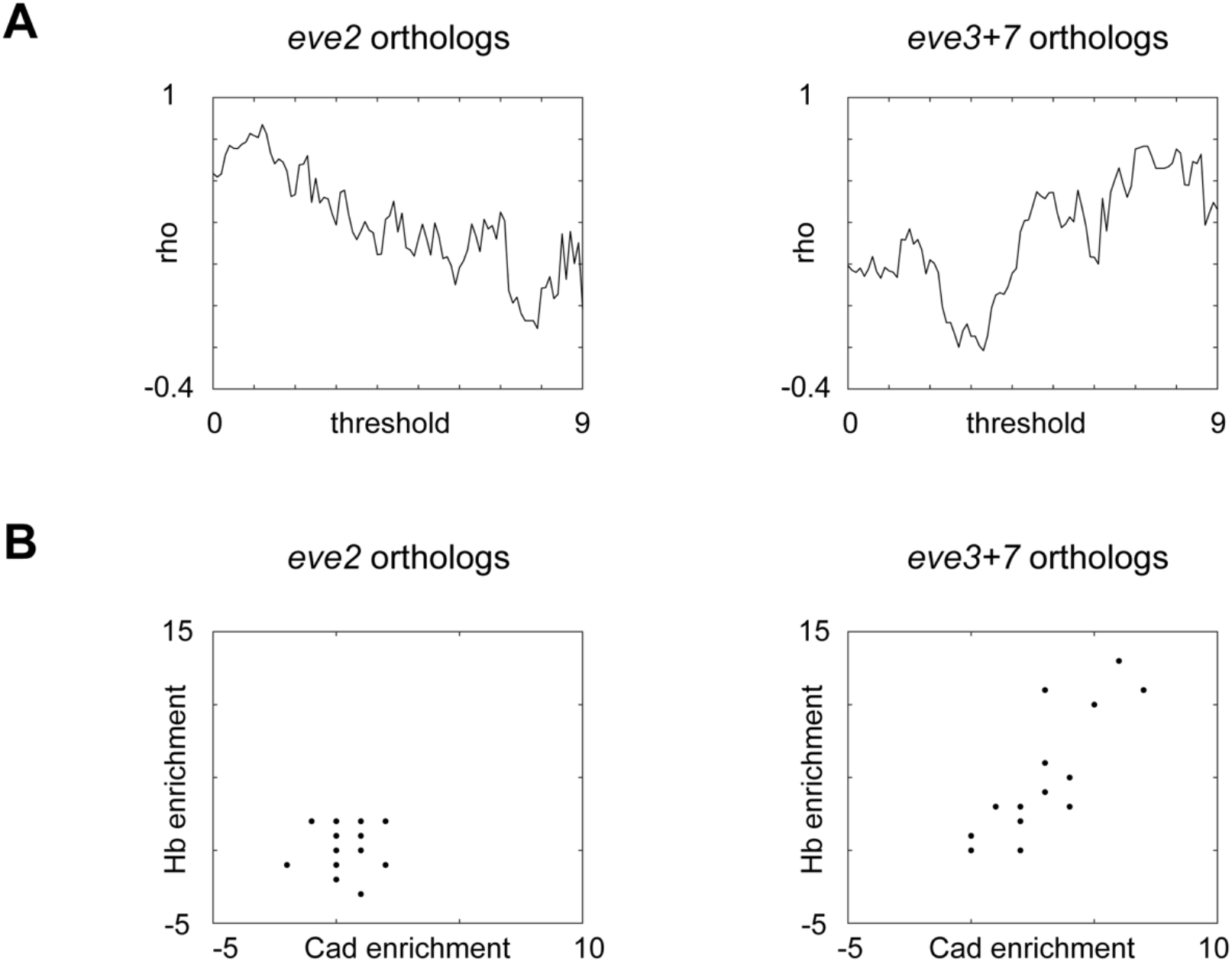
Sensitivity analyses for Caudal and Hunchback enrichment correlations. (A) Spearman correlation for Cad and Hb binding site enrichment is plotted as a function of binding site threshold for *eve2* and *eve3*+*7* orthologs. Binding site threshold refers to the minimum PATSER score for a predicted site to be counted in the analysis. Higher PATSER scores are assumed to reflect higher affinity sites. (B) Hb and Cad enrichment values are plotted for individual *eve2* and *eve3*+*7* orthologs at a binding site threshold of 7.2 – the threshold that maximizes rho in *eve3*+*7*orthologs.

**Fig. S6:**
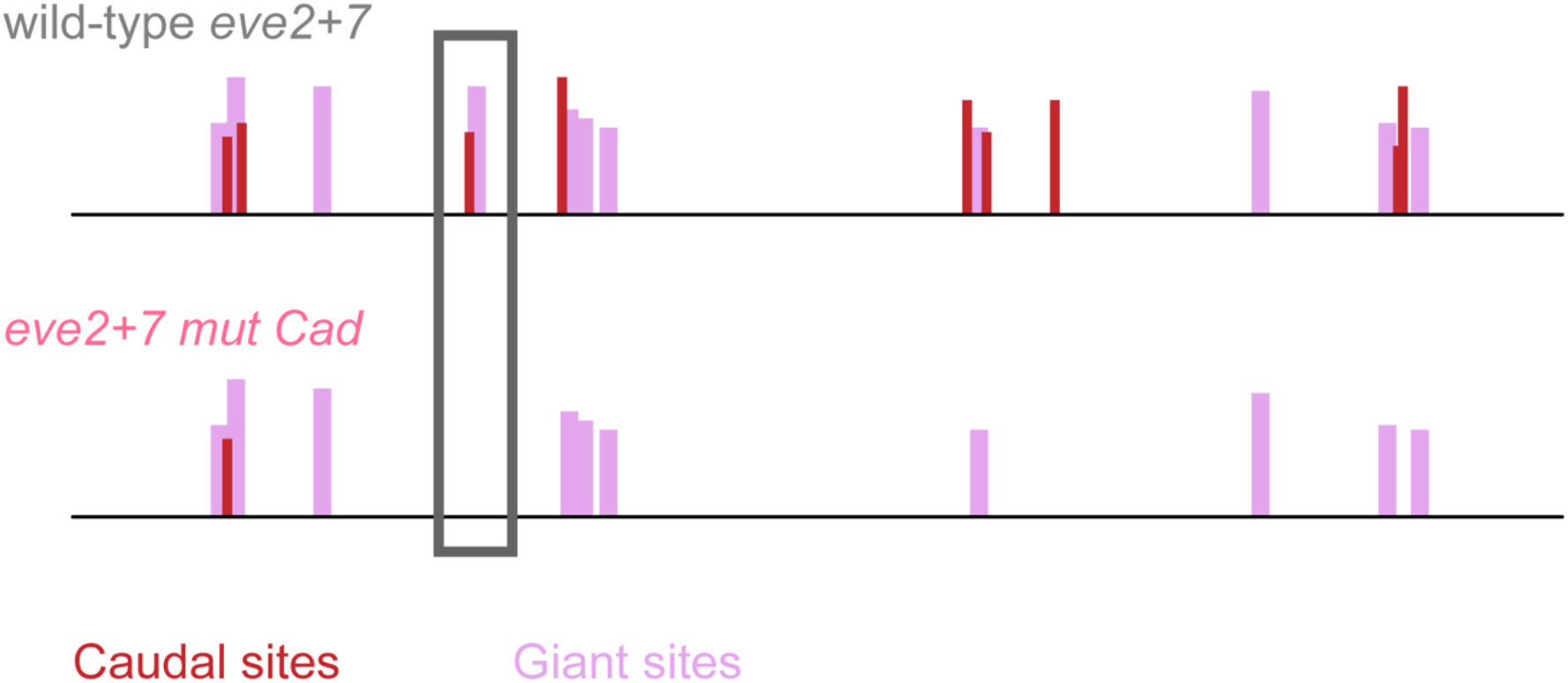
Cad mutations may have disrupted one or more Giant binding sites. Predicted binding for Cad (carmine) and Gt (lilac) are shown in *eve2*+*7* and *eve2+7 mut Cad*. Many predicted Giant binding sites are near predicted Cad sites. One Cad binding site mutation in *eve2+7 mut Cad* (grey box) disrupts a predicted Giant binding site that also overlaps an annotated Giant binding site (47).

